# Identifying condition-related cell-cell communication events using supervised tensor analysis

**DOI:** 10.1101/2023.12.15.571918

**Authors:** Qile Dai, Jingjing Yang, Michael P. Epstein

**Author notes:** Correspondence Authors: M.P.E. and J.Y.

## Abstract

Numerous tools have been developed to infer active cell-cell communication (CCC) events, which are essential for understanding biological processes and diseases. However, existing downstream methods for assessing the relationships between CCC events and biological conditions lack clear interpretation, fail to adjust for confounders, and ignore dependencies among CCC events. To address these limitations, we introduce STACCato, a **S**upervised **T**ensor **A**nalysis tool designed to identify **C**ondition-related **C**ell-cell communic**at**i**o**n events. STACCato employs a tensor-based regression model to enable statistical inference related to the relationships between biological conditions (e.g., disease status, tissue types) and specific CCC events, while adjusting for confounders and CCC dependencies. Through extensive simulations and real-world applications on scRNA-seq datasets of lupus and autism, we demonstrate that STACCato consistently provides improved inference of condition-related CCC events compared to alternative methods. The computational tool implementing the STACCato framework is available on GitHub.

## Introduction

Cell-cell communication (CCC) involves cells exchanging signals to coordinate physiological and developmental functions in multicellular organisms. The study of CCC events, involving interactions between one ligand-receptor pair from a sender cell type to a receiver cell type, is important for elucidating biological processes, exploring disease mechanisms, and inspiring advancements in drug discovery. Using gene expression data profiled by single-cell RNA sequencing (scRNA-seq) technology, multiple computational tools are now available for inferring active CCC events (Jin *et al*., 2021; Efremova *et al*., 2020; Raredon *et al*., 2022; Hu *et al*.; Wang *et al*., 2019; Hou *et al*., 2020; Cabello-Aguilar *et al*., 2020; Armingol *et al*., 2022; Tsuyuzaki *et al*., 2023). Existing CCC inference tools first group individual cells into clusters according to their expression profiles, and then label the cell types of cell clusters based on known marker genes. Next, CCC inference tools evaluate the cell-cell communication strengths by calculating a communication score based on the expression levels of the ligand in a sender cell type and the receptor in a receiver cell type.

Recently, high-throughput sequencing technology advancements have significantly reduced the cost of scRNA-seq, allowing researchers to gather scRNA-seq data from multiple samples under multiple biological conditions (Thompson *et al*., 2022; Perez *et al*., 2022; Nassir *et al*., 2021; Liao *et al*., 2020), such as samples from disease versus healthy control subjects or samples from multiple tissue types. This progress has opened new opportunities for identifying CCC events associated with specific biological conditions. However, existing CCC inference tools primarily focus on estimating the communication strength of CCC events and lack the capacity to rigorously identify CCC events that vary across different biological conditions. One common strategy is to employ an aggregation procedure (Huang *et al*., 2022; Hou *et al*., 2020; Wang *et al*., 2022; Astorkia *et al*., 2022), which first aggregates the scRNA-seq data of all samples within the same condition into one pseudo sample, and then performs CCC inference using the aggregated pseudo sample per condition. CCC events with different communication scores across conditions are identified as condition-related CCC events. Major weaknesses of the aggregation procedure include i) it is not applicable to continuous biological conditions (e.g., age), ii) it disregards heterogeneity among individual samples with the same condition, and iii) the comparison of scores between aggregated pseudo-samples prevents the use of statistical methodology for rigorous testing and inference.

A more recent strategy to identify condition-related CCC events across multiple samples and conditions is to first apply a factor decomposition technique to extract lower-dimensional proxies (i.e., factors) of CCC patterns (Armingol *et al*., 2022; Baghdassarian *et al*., 2023), which can then be compared across conditions. For example, Tensor-cell2cell (Armingol *et al*., 2022) applies factor decomposition to a 4-dimensional communication score tensor (with 4 dimensions corresponding to samples, ligand-receptor pairs, sender cell types, and receiver cell types), and then tests whether the resulting factors are significantly associated with the conditions. Although Tensor-cell2cell allows for statistical inference, its practical limitation is that it tests the condition relatedness of low-dimensional proxies of CCC data rather than the individual CCC events, which complicates interpretation and muddles meaningful biological insights. For instance, in a previous Tensor-cell2cell analysis of a COVID-19 dataset (Armingol *et al*., 2022), after identifying factors significantly associated with COVID-19, the interactions between the ligand MIF and the receptors CD74 and CD44 were found among the top five most active ligand-receptor pairs in two significant factors but with opposite associations. That is, one factor was positively associated with COVID-19, suggesting a detrimental role for MIF-CD74/CD44, while the other was negatively associated, indicating a protective role. Such scenarios make it impossible to conclude the overall relationship between CCC events and the condition of interest and limit the practical utility of Tensor-cell2cell such as for guiding therapeutic strategies or prioritizing specific interactions.

An additional practical limitation of both the aggregation strategy and the factor decomposition strategy for studying condition-related CCC events is that neither account for other important sample-level variables (e.g., processing batch, age, gender, and ancestry) that can confound the association between CCC events and biological conditions of interest. Failing to account for potential confounding variables may mask true biological associations between CCC events and biological conditions. Even more concerning, lack of confounder adjustment can lead to false positive associations that could result in misguided interpretations. There is a clear need for improved statistical methodology for condition-related CCC analysis that can adjust for sample-level confounders to yield more rigorous inference.

To address limitations of existing procedures for detecting condition-related CCC events, we introduce the **S**upervised **T**ensor **A**nalysis tool for identifying **C**ondition-related **C**ell-cell communic**at**i**o**n (**STACCato**). STACCato is a tensor-based regression model, which takes the 4-dimensional communication score tensor (with 4 dimensions corresponding to samples, ligand-receptor pairs sender cell types, and receiver cell types) as the outcome response variable, and the biological conditions and other sample-level covariates as independent variables (**Fig. 1**). The communication score tensor can be generated using any available CCC inference tool. Unlike existing procedures for detecting condition-related CCC events, STACCato makes inference directly on the relation between CCC events and the biological condition of interest, while accounting for sample heterogeneity and confounding variables. Unlike traditional regression models that test each CCC event separately, STACCato considers all CCC events simultaneously which is preferred because of the interdependency among CCC events (e.g., the interactions of the same ligand-receptor pair across different sender and receiver cell types are dependent). Here, we show that STACCato’s ability to account for dependency of CCC events leads to improved inference over the use of traditional regression models. Thus, we feel STACCato provides a valuable and flexible downstream tool for identifying condition-related CCC events in multi-sample, multi-condition scRNA-seq datasets.

**Fig. 1.**
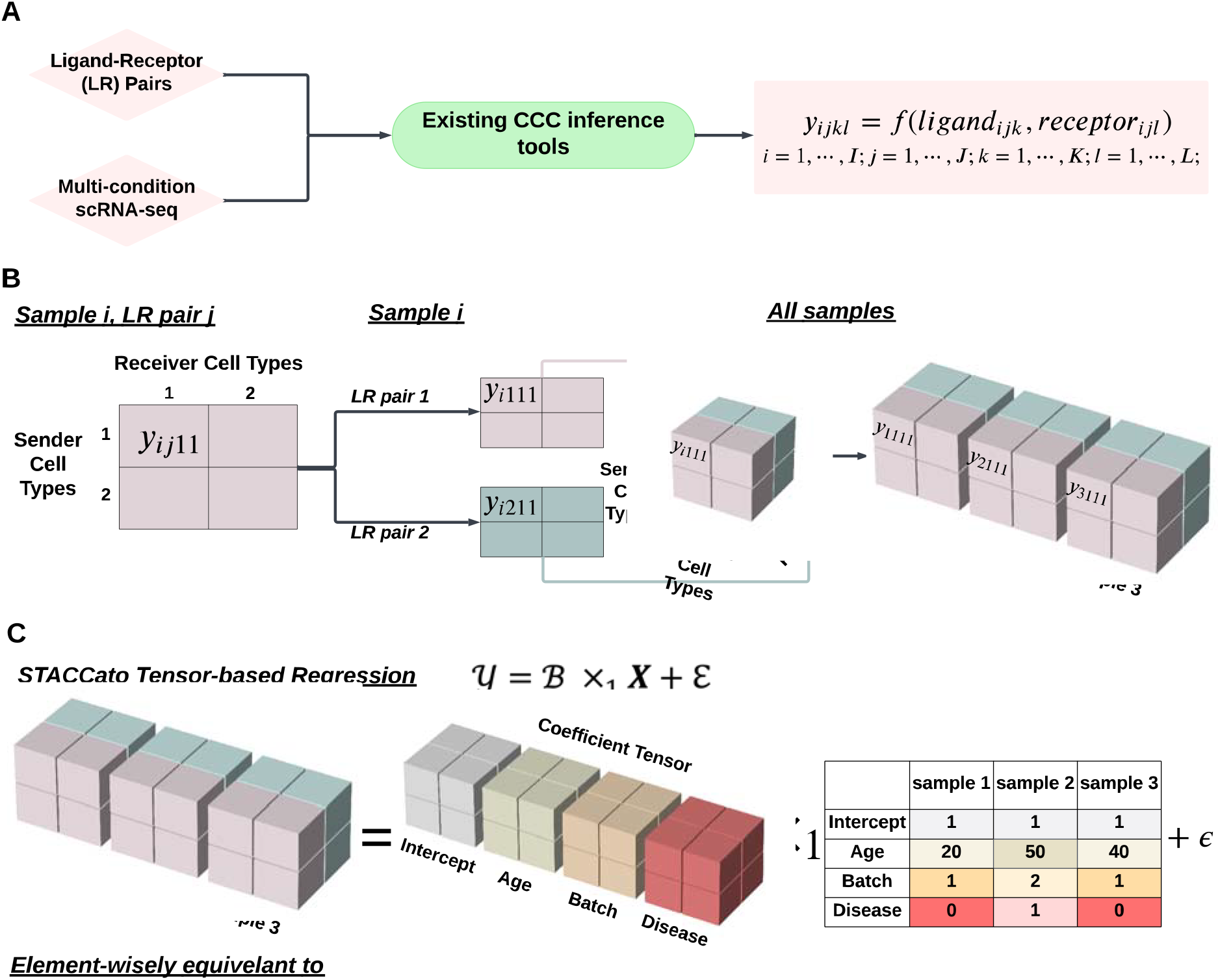
STACCato framework. **(A)**. Given the scRNA-seq data of sample *i* and a ligand-receptor pair *j*, cell-cell communication (CCC) score is given by a function of the expression levels of ligand in sender cell type *k* (*ligand*_*ijk*_) and receptor in receiver cell type *l* (*receptor*_*ijl*_); **(B)**. Given the scRNA-seq data of sample *i*, CCC scores are calculated for ligand-receptor pair *j* across all sender and receiver cell types. CCC scores are then organized into a communication score matrix with sender cell types as rows and receiver cell types as columns. For each sample *i*, communication score matrices are repeatedly calculated for all ligand-receptor pairs and organized into a 3-dimensional communication score tensor. The 3-dimensional communication score tensors are repeatedly constructed for all samples and then combined into a 4-dimensional communication score tensor; **(C)**. STACCato then uses a tensor-based regression to estimate the coefficient tensor representing the effects of sample-level variables on CCC events. While this example tensor contains only 2 cell types and 2 ligand-receptor pairs, the framework is generalizable to any number of cell types and ligand-receptor pairs.

In subsequent sections, we first introduce the analytical framework of STACCato. We then illustrate STACCato with application to two real datasets: the Systemic Lupus Erythematosus (SLE) scRNA-seq dataset (Thompson *et al*., 2022; Perez *et al*., 2022) of peripheral blood mononuclear cells (PBMC) samples from 145 managed SLE patients and 48 healthy controls, and the Autism Spectrum Disorder (ASD) scRNA-seq dataset (Nassir *et al*., 2021) consisting of prefrontal cortex (PFC) samples from 13 ASD patients and 10 controls. We further demonstrate the features and advantages of STACCato through extensive simulations conducted under various study designs. Finally, we conclude with a discussion.

## Methods

### Overview of STACCato framework

For each sample in the scRNA-seq dataset, STACCato first uses existing CCC inference tools to estimate the communication scores of each CCC event involving the interaction of one ligand-receptor pair from one sender cell type to one receiver cell type (**Fig. 1A**). STACCato then generates a 4-dimensional communication score tensor with four dimensions representing samples, ligand-receptor pairs, sender cell types, and receiver cell types (**Fig. 1B**). Next, STACCato employs a tensor-based regression method that takes the 4-dimensional communication score tensor as the outcome response variable and sample-level variables including the biological condition of interest as independent variables. That is, STACCato incorporates sample-level information (such as batches, age, gender, and the biological condition of interest) to estimate a coefficient tensor containing the effects of sample-level variables on CCC events (**Fig. 1C**). Finally, STACCato conducts parametric bootstrapping to rigorously assess the significance of these estimated coefficients. We describe this tensor-based regression framework as follows.

### Generation of communication scores

STACCato estimates the communication scores of CCC events using existing inference tools. Such tools leverage scRNA-seq expression data to quantify communication scores among various ligand-receptor pairs procured from literature-curated lists in each pair of sender-receiver cell types (**Fig. 1A**). The following communication score *y*_*ijkl*_ is generated for the CCC event in sample involving the interaction of ligand-receptor pair *j* from sender cell type *k* to receiver cell type *l*

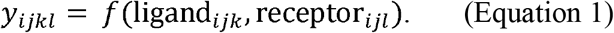

Here, ligand_*ijk*_ denotes the expression of the ligand of the ligand-receptor pair *j* in sender cell type *k* of sample ; receptor_*ijl*_ denotes the expression of the receptor of the ligand-receptor pair *j* in receiver cell type of sample ; and *f* denotes the scoring function. In this study, we utilized the product scoring function *y*_*ijkl*_ ligand_*ijk*_ × receptori_*ijl*_ as used by Connectome (Raredon *et al*., 2022) and NATMI (Hou *et al*., 2020), along with the consensus list of ligand-receptor pairs provided in the LIANA+ package (Dimitrov *et al*., 2022, 2023) (see Data and materials availability). One can also employ different scoring functions as used by different existing inference tools and different lists of literature-curated ligand-receptor pairs.

### Regression models for identifying condition-related CCC events

Assume communication scores (*y*_*ijkl*_ as in Equation 1) have been generated from a multi-sample scRNA-seq dataset of *I* samples, with *J* ligand-receptor pairs, *K* sender cell types, and *L* receiver cell types. Assume *Q* sample-level variables (*x*_*i*,1_, *x*_*i*,2_,…,*x*_*i,Q*_) are collected for each sample, including the biological condition of interest (e.g., disease/normal) and additional sample covariates (e.g., batch, age, and gender). Condition-related CCC events can be identified by using the following covariate-adjusted regression models:

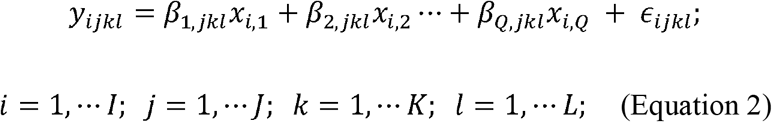

where *ϵ*_*ijkl*_ ∼ *N* (0, *σ*^*2*^)denotes random error that follows a Gaussian distribution with mean 0 and standard deviation σ. The above regression model (Equation 2) aims to estimate the coefficients {*β*_*q,jkl*_ *q*= 1,…,*Q*}, which denote the effects of sample-level variables on the communication score of the CCC event (*y*_.*jkl*_) involving the interaction of ligand-receptor pair *j* from sender cell type *k* to receiver cell type *l*. For example, for a scRNA-seq dataset with samples collected from disease and healthy control subjects, we can set *x*_*i,Disease*_ = 0 for healthy controls and *x*_*i,Disease*_ = 1 for disease subjects. Then CCC events with significant *β*_*Disease,jkl*_ are the CCC events affected by the disease. The larger the magnitude of the coefficient, the greater the difference in communication magnitudes between the disease and control conditions. We refer to CCC events with *β*_*Disease,jkl*_ significantly different from 0 as CCC events related to the disease condition of interest, while adjusting for additional confounding covariates such as batches, age, and gender.

A straightforward way to estimate *β*_*q,jkl*_ in Equation 2 is through the separate regression approach, by fitting the regression models independently for each CCC event associated with ligand-receptor pair *j*, sender cell type *k*, and receiver cell type *l*. However, this separate regression approach ignores the dependencies among CCC events. For example, the interactions of the same ligand-receptor pair *j* across different sender and receiver cell types are known to be dependent, and thus *β*_*q,jkl*_ is dependent of *β*_*q,jk′l′*_ with *k* ≠ *k*’ and *l* ≠ *l*’.

### Tensor-based regression for joint inference of CCC events

To account for interdependency among all CCC events, STACCato employs a tensor-based regression method to jointly estimate ***β***_*jkl*_ *=*[β,*jkl*,…, *β*_*Q,jkl*_]^*T*^ for all *j* = 1,…*J* ; *k* = 1,…*K*; *l* = 1,… *L*. First, STACCato arranges the communication scores into a 4-dimensional communication score tensor *y* ∈ ℝ^*I×J×K×L*^, with the (*i,j,k,l*) entry corresponding to *y*_*ijkl*_ in Equation 2 (see details about communication score tensor construction in Supplementary Methods). **Fig. 1B** shows a graphic example of building the 4-dimensional communication score tensor for a scRNA-seq dataset with three samples, two ligand-receptor pairs, and two sender-receiver cell types. Then the regression models in Equation 2 can be written in a tensor format as follows:

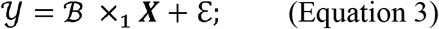

Where **x** ∈ ℝ^*i × Q*^ denotes sample-level design matrix of *Q* variables; ×_1_ denotes multiplying a tensor by a matrix in the tensor’s first dimension; ε ∈ ℝ^*I×J×K×L*^ denotes a 4-dimensional tensor with the (*i,j,k,l*) entry corresponding to *ϵ*_*ijkl*_ in Equation 2. **Fig. 1C** shows a graphic example representation of the tensor-based regression model in Equation 3 for three samples, with disease, age, and batch as sample-level variables.

STACCato aims to estimate the coefficient tensor ℬ ∈ ℝ^*QXJXKXL*^, with the (*q,j,k,l*) entry corresponding to *β*_*q,jkl*_ in Equation 2. According to the tensor decomposition technique (Hu *et al*., 2022), ℬ can be viewed as a core tensor 𝒢 multiplied by 4 factor matrices ***M***_*Q*,_ ***M***_*j*,_ ***M***_*K*,_ ***M***_*L*_,

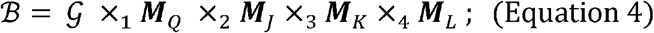

where ×_*d*,_ *d* = 1,2,3,4 denotes multiplying a tensor by a matrix in the tensor’s *d*th dimension. For the convenience of presentation, we use 𝒢 × {***M***_*Q*_, ***M***_*j*_, ***M***_*K*_, ***M***_*L*_} to denote the tensor-by-matrix product in Equation 4. Then the full STACCato model is given by:

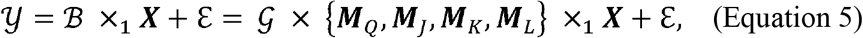

where 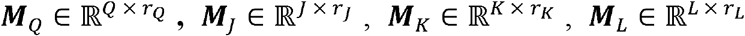 are factor matrices, 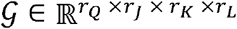 denotes the core tensor whose entries show the level of interaction among the factors from different dimensions, and (*r*_*Q*_, *r*_*j*_, *r*_*K*_, *r*_*L*_) denote the decomposition ranks of each factor matrix (see details about rank determination in Supplementary Methods). These factor matrices have orthonormal columns (i.e., factors), which can be thought of as the principal components for each dimension. STACCato uses the QR-adjusted optimization algorithm proposed by Hu et al. (Hu *et al*., 2022) to estimat ℬ, 𝒢, ***M***_*Q*_, ***M***_*J*_, ***M***_*K*_ ***M***_*L*_. The significance levels of estimated coefficients in ℬ are assessed by parametric bootstrap(Efron and Tibshirani, 1994). We describe the details of QR-adjusted optimization algorithm and bootstrap procedure in Supplementary Methods.

### Accounting for dependencies among all CCC events

Based on the tensor-based regression model described above, STACCato jointly estimates all coefficients in ℬ and naturally accounts for the dependencies among all CCC events. For example, based on Equation 5, *β*_*q,jkl*_ in the coefficient tensor ℬ that quantifies the effect of covariate *q* on the CCC event with ligand-receptor pair *j*, sender cell types *k*, and receiver cell type is given by

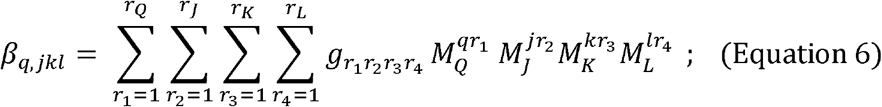

where 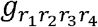 denotes the (*r* _1_, *r*_2_, *r*_*3*_, *r*_4_) entry of 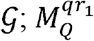 denotes the entry in the *q*^*th*^ row and 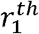 column of *M*_*Q*_ ; and 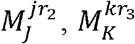, and 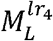 are similarly defined. Likewise, the coefficient *β*_*q,jk’l’*_ of covariate *q* on another CCC event with the same ligand-receptor pair *j*, but different sender cell types *k*’ ≠*k* and receiver cell type *l*^′^ ≠ *l* is given by

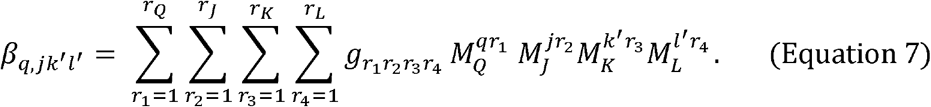

We can see that the estimates of *β*_*q,jkl*_ in Equation 6 and *β*_*q,jk’l’*_ in Equation 7 involve the same 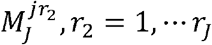, related to the same ligand-receptor pair *j*; thus naturally accounting for the dependencies between these two CCC events arising from sharing the same ligand-receptor pair *j*. Similarly, for CCC events with the same sender cell type *k*, the corresponding coefficient estimates share the same 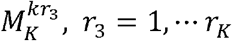; for CCC events with the same receiver cell type *l*, the corresponding coefficient estimates share the same 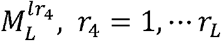. Therefore, through these shared factors for estimating CCC events involving shared components of ligand-receptor pairs, sender cell types, and receiver cell types, STACCato effectively accounts for the complex dependencies among all CCC events.

## Results

### Applying STACCato to identify CCC events associated with SLE

We applied STACCato to a scRNA-seq dataset of PBMC samples from 145 managed SLE subjects and 48 healthy controls (Thompson *et al*., 2022; Perez *et al*., 2022) to identify CCC events associated with SLE while adjusting for age, gender, self-reported ancestry, and processing batches (see Supplementary Methods for details). We considered 9 cell types present in over 80% of the samples: B cells, natural killer cells (NK), proliferating T and NK cells (Prolif), CD4^+^ T cells, CD8^+^ T cells, CD14^+^ classical monocytes (cM), CD16^+^ nonclassical monocytes (ncM), conventional dendritic cells (cDC), and plasmacytoid dendritic cells (pDC).

First, we calculated the communication scores for 362 ligand-receptor pairs from the LIANA+ consensus list that are present in all samples within all pairs of 9 sender and receiver cell types. Second, we generated the 4-dimensional communication score tensor with dimensions of 193 × 362 × 9 × 9, for analyzing 193 samples, 362 ligand-receptor pairs, 9 sender cell types, and 9 receiver cell types, with 95.9% of the communication scores in the tensor are zeros. Third, by using the rank selection strategy as described in the Supplementary Methods, we selected the decomposition rank ***r*** = (*r*_*Q*_ =6, *r*_*j*_ = 9, *r*_*K*_ = 3, *r*_*L*_ = 7) for STACCato to estimate the coefficient tensor. Last, we conducted 4,999 iterations of bootstrapping resampling to assess the significance levels of the estimated SLE disease effects.

We identified disease effects with bootstrapping p-values < 0.05 and estimated magnitudes > 0.02 as significant disease effects (**Fig. S1**). We identified a total of 1,659 CCC events that were positively associated with SLE, considerably more than the 70 CCC events that were negatively associated (**Fig. S2**). These positive associations are evenly distributed across various cell types. In contrast, among the negative associations, cM predominantly acts as the major sender cell type and NK is the primary receiver cell type.

**Fig. 2** displays the top 1,000 significant SLE-related CCC events with top magnitudes for their estimated disease effects. We identified positive associations involve multiple HLA Class I genes (including HLA-A, HLA-B, HLA-C, and HLA-E) and Class II genes (including HLA-DRB1, HLA-DRA, and HLA-DPB1). HLA genes are strongly associated with autoimmune diseases because of their crucial roles in presenting self-antigens to the immune system, which can lead to an inappropriate immune response and autoantibody production when dysregulated (Molineros *et al*., 2019; Raj *et al*., 2016). Significant SLE-related CCC events involving the S100 family proteins, particularly S100A8 and S100A9, were also identified across various cell types. In particular, the CCC events between S100 family proteins and the Cluster of Differentiation 68 (CD68) are positively associated with SLE. Previous studies show that S100A8 and S100A9 tightly bind to CD68 and that their interaction could regulate the cells’ immune functions in rat (Okada *et al*., 2016).

**Fig. 2.**
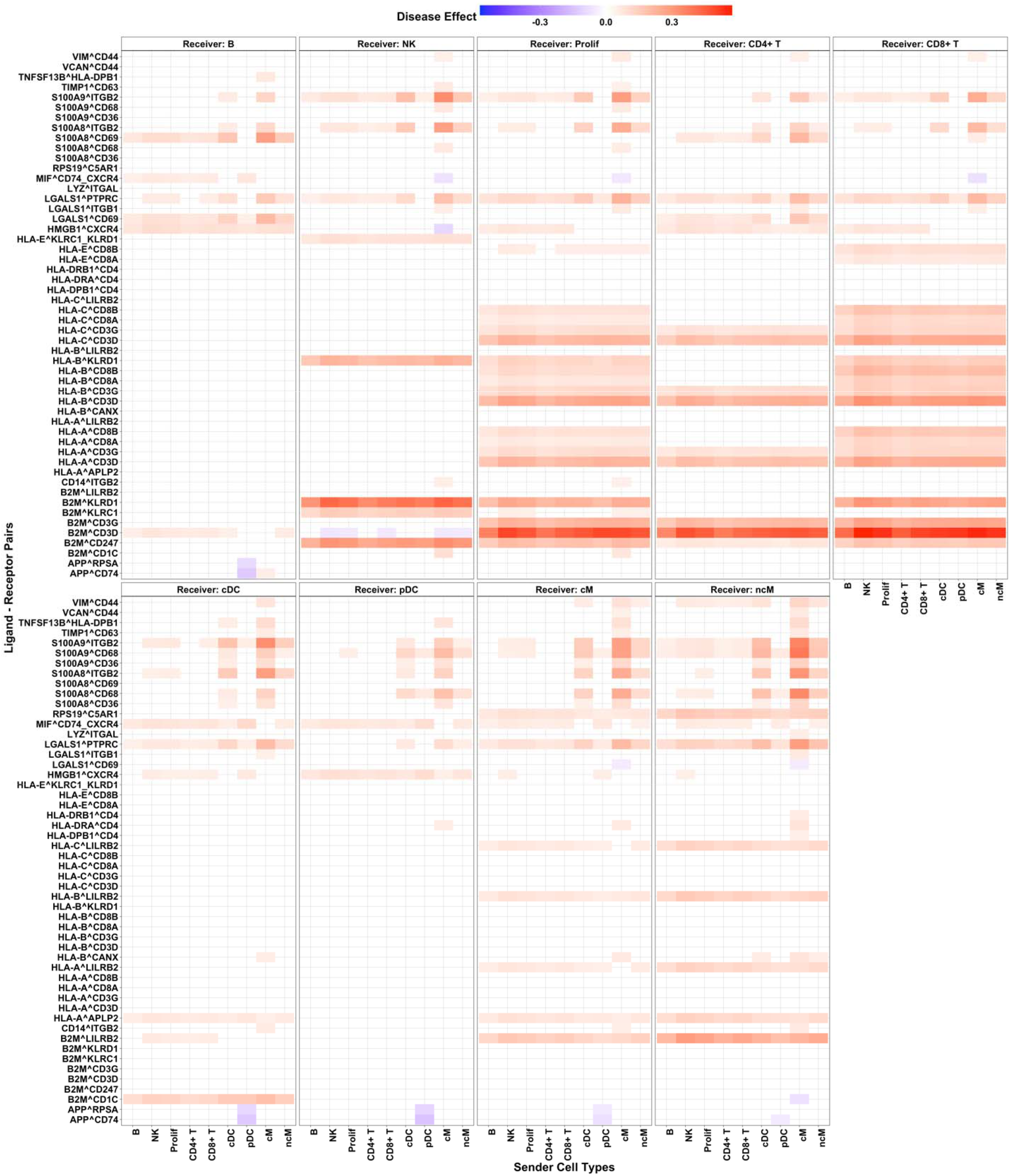
STACCato results with the SLE dataset. Top 1000 significant communication events with top magnitudes of their estimated SLE disease effects. Positive disease effects colored in red indicate positive associations between CCC events and SLE, while negative disease effects colored in blue indicate negative associations. Each panel represents a different receiver cell type indicated in the panel title. Within each panel, each grid shows the estimated SLE disease effect for a specific CCC event, with involved ligand-receptor pair labeled on the y-axis, and the sender cell types indicated on the x-axis.

### Applying STACCato to identify CCC events associated with ASD

We applied STACCato to the snRNA-seq dataset of postmortem tissue samples of prefrontal cortex from 13 ASD patients and 10 controls (Nassir *et al*., 2021) to identify CCC events associated with ASD, while adjusting for age, gender, and processing batches (see Supplementary Methods for detailed descriptions about the dataset). We considered 17 sender/receiver cell types present in over 80% of the samples: fibrous astrocytes (AST-FB), protoplasmic astrocytes (AST-PP), Endothelial, parvalbumin interneurons (IN-PV), somatostatin interneurons (IN-SST), SV2C interneurons (IN-SV2C), VIP interneurons (IN-VIP), layer 2/3 excitatory neurons (L2/3), layer 4 excitatory neurons (L4), layer 5/6 corticofugal projection neurons (L5/6), layer 5/6 cortico-cortical projection neurons (L5/6-CC), Microglia, maturing neurons (Neu-mat), NRGN-expressing neurons (Neu-NRGN-I), NRGN-expressing neurons (Neu-NRGN-II), Oligodendrocyte precursor cells (OPC), and oligodendrocytes.

Similarly, we first calculated the communication scores for 2610 ligand-receptor pairs from the LIANA+ consensus list that were present in all samples, and then generated the 4-dimensional communication score tensor with dimensions of 23 × 2610 × 17 × 17 for the CCC events of 23 subjects, 2610 ligand-receptor pairs, 17 sender cell types, and 17 receiver cell types, with 98.1% of the communication scores in the tensor are zeros. Next, we selected the decomposition rank ***r*** = (*r*_*Q*_ =5, *r*_*j*_ *=* 10, *r*_*K*_ = 6, *r*_*L*_ = 7) for estimating the coefficient tensor, and conducted 4,999 iterations of bootstrapping resampling to assess the significance levels of the estimated ASD disease effects. As a result, we identified 8,347 CCC events that were significantly associated with ASD, with p-values < 0.05 and magnitudes of estimated effects > 0.02 (**Fig. S3**). In **Fig. 3**, we display top 1,000 significant ASD-related CCC events with top magnitudes of estimated disease effects. We identified strong associations between ASD and CCC events involving Neurexins (NRXN) and Neuroligins (NLGN), which are crucial for synapse formation and function(Südhof, 2008). NRXN1 mutations have been linked to synaptic dysfunction, representing a major underlying mechanism for ASD (Guang *et al*., 2018). Additionally, the Calcium Signaling Pathway, involving genes of *CALM1, CALM3, CACNA1C, GRM5, PDE1A*, and *RYR2*, showed negative associations with ASD (**Fig. 3**). Dysregulated calcium signaling has been linked to several neurodevelopmental disorders, including ASD, where it can lead to deficits in neuronal communication and synaptic function (Splawski *et al*., 2004).

**Fig. 3.**
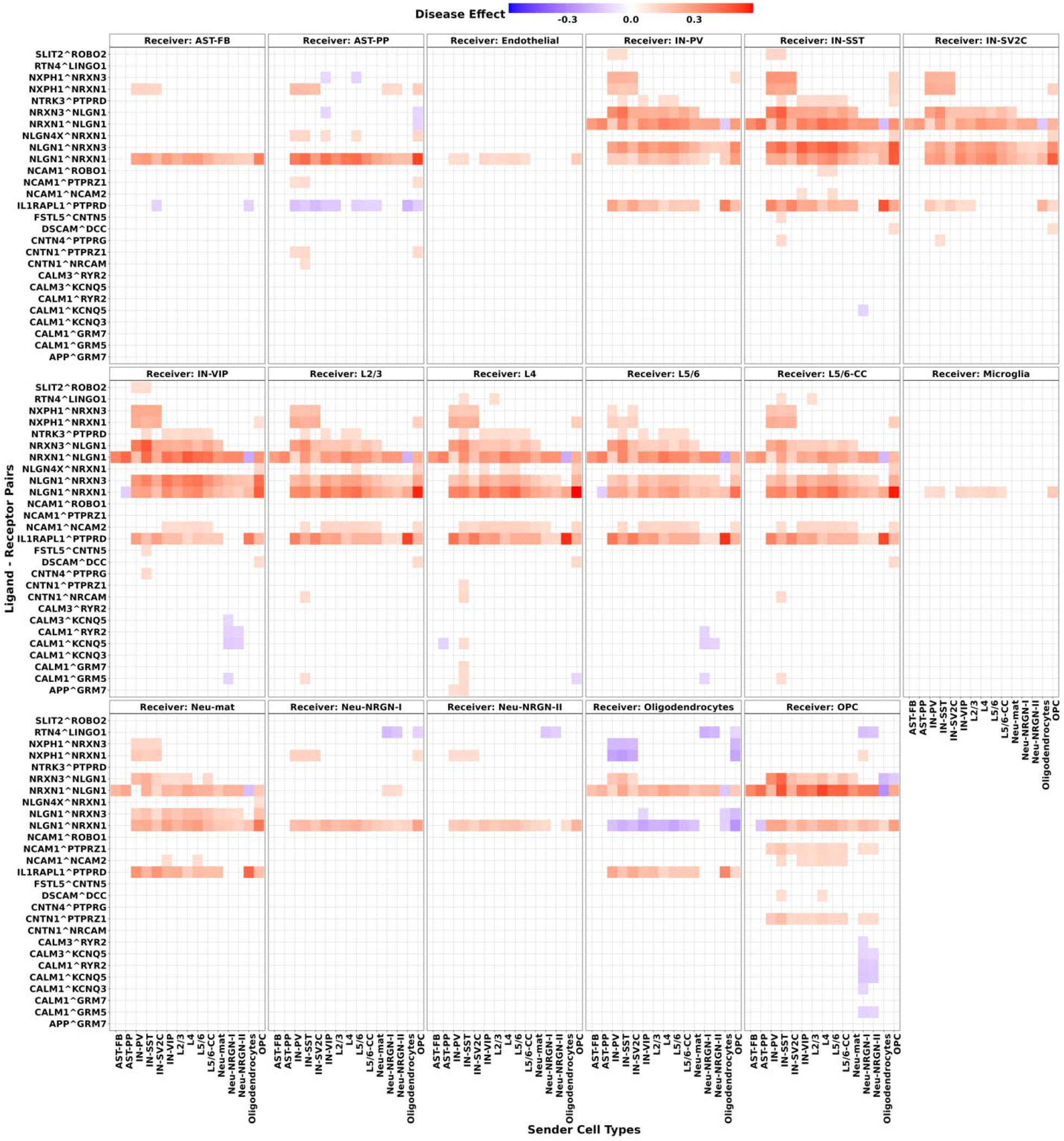
STACCato results with the ASD dataset. Top 1000 significant communication events with top magnitudes of their estimated ASD disease effects. Positive disease effects colored in red indicate positive associations between CCC events and ASD, while negative disease effects colored in blue indicate negative associations. Each panel represents a different receiver cell type indicated in the panel title. Within each panel, each grid shows the estimated ASD disease effect for a specific CCC event, with involved ligand-receptor pair labeled on the y-axis, and the sender cell types indicated on the x-axis.

### Evaluating the impact of confounding variables for identifying condition-related CCC events

One noteworthy aspect of the SLE dataset is its highly unbalanced study design (**Table S1**), particularly in Batch 1, where 96.33% (105 out of 109) of the subjects are SLE patients, and only 3.67% (4 out of 109) are healthy controls. Consequently, batch effects confounded the associations between CCC events and SLE disease condition. To evaluate the impact of confounding variables on the identification of condition-related CCC events, we applied STACCato to the SLE dataset with three distinct models, each incorporating different sample-level variables: Model 1 that considers all available sample-level variables of disease status, batches, and all other available covariates including age, gender, and ancestry (whose results were shown in **Fig. 2**); Model 2 that considers disease status and batches only; and Model 3 considers disease status only.

We observed substantial changes in the estimated disease effects before batch adjustment (Model 3) and after adjustment (Model 2 and Model 1). We show the estimated disease effects for the top 40 CCC events with the largest changes before and after adjusting for batch effects in **Fig. S4**. For example, the estimated disease effect on the CCC event between APP and CD74 in pDC cells is 0.14 in Model 3, before adjusting for batch effects. After adjusting for batch in Model 2 and Model 1, the estimated disease effect of this CCC event is of similar magnitude but in opposite direction (−0.14 and -0.15, respectively). By examining the communication scores of this CCC event, we observed a significant batch effect, with communication scores being larger in Batch 1 compared to Batch 2 (**Fig. S5A**). Given that 96.33% of the subjects in Batch 1 are SLE patients, this batch effect can create a misleading impression that SLE patients have larger or comparable communication scores compared to healthy controls (**Fig. S5B**). Instead, after adjusting for batch effects, the scores in SLE patients are actually lower than those in healthy controls within both batches (**Fig. S5C**), consistent with the disease effects estimated in Model 1 and Model 2. These findings underscore how confounding variables can distort true associations and emphasize the importance of considering confounding variables in identifying condition-related CCC events.

We also examined the impact of batch information on our ASD results by fitting three distinct STACCato models with Model 1 considering disease status and all available covariates including batches, age, and gender (with results shown in **Fig. 3**), Model 2 considering disease status and batches only, and Model 3 considering disease status only. Unlike the SLE dataset, the ASD dataset exhibits a fairly balanced design (**Table S2**). Consequently, batch is no longer a confounding factor. As anticipated, the estimated disease effects before and after adjusting for batch effects are consistent (**Fig. S6**).

### Comparing STACCato to the separate regression approach

We compared STACCato to the separate regression approach (which ignores interdependency among CCC events) by analyzing the consistency of results between the full dataset and sub-datasets containing 70% of the samples selected randomly without replacement from the full dataset. We created 20 sub-datasets and calculated two metrics for each sub-dataset and each method: (1) the Pearson correlation coefficient to compare the estimated disease effects in the sub-dataset with those in the full dataset, and (2) the proportion of the top 1000 significant CCC events (those with the largest magnitudes of disease effects) in the full dataset that were also detected in the sub-dataset. We conducted paired t-test for each of the metrics to test the significant differences between STACCato and separate regression methods.

STACCato consistently yields more robust inference than the separate regression approach, with significantly higher correlation coefficients and greater overlap proportions in both the SLE (**Fig. S7**) and ASD datasets (**Fig. S8**), with t-test p-values < 0.01. This is likely because the separate regression approach is more sensitive to noise and outliers specific to each event, leading to potentially less reliable and more variable outcomes in the sub-datasets. In contrast, the results given by STACCato are more stable and more robust to noise, due to joint inference on all CCC events using the tensor-based regression model. Through joint inference, information can be borrowed across correlated CCC events, thus resulting in more robust and stable estimates for the disease effects on CCC events.

### Simulation studies based on the SLE dataset

To further investigate how sample-level variables affect the identification of condition-related CCC events, we conducted in-depth simulation studies with various study designs. We simulated communication score tensors 𝒴 ∈ ℝ^*I*×*J*×*K*×*L*^ for 60 subjects (*I*=60) from the tensor-based regression model as in Equations 5, taking the estimated core tensor of 𝒢, and factor matrices ***M***_*Q*_, ***M***_*J*_, ***M***_*K*_, ***M***_*L*_ from the SLE dataset as the truth. We simulated the design matrix x with an intercept column and the covariates of disease status and batch variables. The elements of εwere independently simulated from a normal distribution with mean 0 and three different standard deviations (σ =0.05,0.08,0.1) to mimic varying levels of noises, where σ = 0.05 was the standard error of the estimation residuals from the SLE data. We simulated the data for 30 disease subjects and 30 healthy controls processed in two batches. We considered three study designs: (1) balanced design with 15 controls and 15 disease subjects in each of these two batches; (2) moderate unbalanced design with 20 controls and 10 disease subjects in Batch 1, and 10 controls and 20 disease subjects in Batch 2; (3) extreme unbalanced design with 30 controls and 5 disease subjects in Batch 1, and 0 controls and 25 disease subjects in Batch 2.

We applied STACCato to the simulated data of each simulation scenario with two models: Model 1 considers disease status and batch variables, and Model 2 considers only disease status. We also compared to the separate regression model that ignored CCC event dependencies. We calculated the mean squared errors (MSEs) of the estimated disease effects and the standard deviation of estimation errors by each model across 100 simulations, for each simulation scenario. **Fig. 4** illustrates that, for simulation data generated under all levels of noises, neglecting the batch confounder by STACCato in both moderate and extreme unbalanced designs leads to elevated MSEs (green vs. red bars), with MSEs increasing dramatically as the degree of imbalance became more extreme. Although both STACCato Model 1 and separate regression model account for the confounding batch covariate, STACCato obtained lower MSEs across all considered simulation scenarios (red vs. blue bars), highlighting the advantage of using the tensor-based regression method to account for correlations among CCC events. Additionally, the MSEs by separate regression model increased dramatically when the level of noises increased, while the MSEs by STACCato Model 1 remain approximately the same. These simulation results (which are consistent with our real-data findings) demonstrate the importance of considering potential confounding covariates when conducting condition-related CCC inference and the value of accounting for CCC dependencies using the tensor regression model employed by STACCato.

**Fig. 4.**
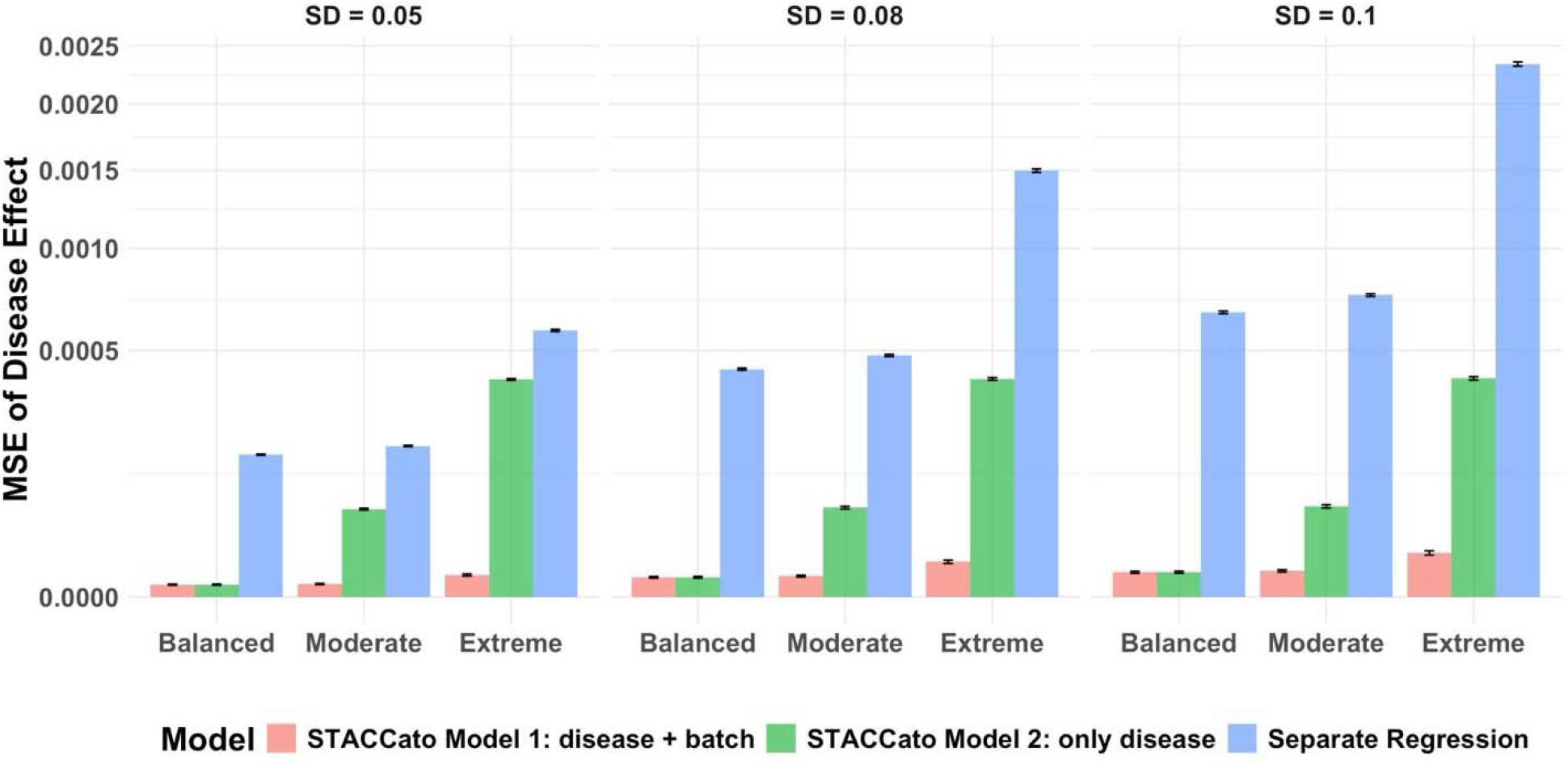
Simulation study results by two comparison models of STACCato and the separate regression model. Simulation data were generated with three different standard deviations (*σ* = 0.05, 0.08, 0.1) to mimic varying levels of noises, as shown in three panels. Bar plots were made for the mean squared errors (MSEs) of the estimated disease effects across 100 simulations in each of the balanced, moderate unbalanced, and extreme unbalanced simulation scenarios, by each model. MSEs by Model 1 considering disease status and batch were plotted in red bars; MSEs by Model 2 considering disease status only were plotted in green bars; and MSEs by the separate regression considering disease status and batch were plotted in blue bars. The standard deviations of the estimation errors across 100 simulations were shown on top of the bars.

### Computational efficiency of STACCato

While fitting of a single STACCato tensor-based regression only takes seconds, assessing the significance level of estimated effects by bootstrapping (requiring multiple iterations of model fitting) takes hours of CPU time. We conducted the computational benchmarks using one Intel(R) Xeon(R) processor (2.10 GHz). For a simulated dataset of 100 samples, 10 sender and receiver cell types, 600 ligand-receptor pairs, and 10 sample-level variables, 99 iterations of bootstrap resampling took around 11 minutes and ∼1.3 GB memory usage on the upper-bound.

Since the numbers of cell types and sample-level variables generally do not vary much in practice, we investigated how the computation CPU time and upper-bound memory usage vary with the number of samples and the number of ligand-receptor pairs, for 99 iterations of bootstrapping resampling. We simulated datasets with 10 sender and receiver cell types and 10 sample-level covariates, and various numbers of samples (*I* = 25, 50, 75, 100) and ligand-receptor pairs (*J* = 150, 300, 450, 600). We demonstrated that computational time increased approximately linearly with the number of samples (**Fig. S9A**) and quadratically with the number of ligand-receptor pairs (**Fig. S9C**). The upper bound memory usage changed approximately linearly with the numbers of samples and ligand-receptor pairs (**Fig. S9B** and **Fig. S9D**).

## Discussion

We present STACCato, a computational tool that utilizes multi-sample multi-condition scRNA-seq data to identify CCC events associated with biological conditions (e.g., disease status, multiple time points, different tissue types). STACCato utilizes a tensor-based regression model to jointly estimate the influence of the condition of interest on CCC events, while adjusting for potential confounding variables and incorporating dependencies among all CCC events. We applied STACCato to analyze a real SLE scRNAseq dataset with an extremely unbalanced design (Perez *et al*., 2022; Thompson *et al*., 2022) and a real ASD scRNAseq dataset with a balanced design (Nassir *et al*., 2021). Additionally, we conducted in-depth simulation studies to mimic real data with balanced and unbalanced study designs and different noise levels. Our real data application and simulation results demonstrate STACCato’s capability to account for available sample-level confounding variables and dependencies among CCC events, thereby enabling more accurate and robust inferences of the associations between CCC events and the biological condition of interest. While our application and simulation studies primarily focused on identifying disease-related CCC events, STACCato can also be applied to other complex biological conditions, such as various ages, multiple tissue types, and multiple time points, allowing for a broader range of investigative possibilities.

In practice, a common approach to address confounding batch effects in scRNA-seq data analysis is to use the data after adjusting for batch effects for downstream analyses. However, as noted by Nygaard et al. (Nygaard *et al*., 2016), batch-effect removal tools typically eliminate the point estimates of these effects while overlooked the estimation errors of these effects. As a result, even if the point estimates of the original batch effects are eliminated, the estimation errors may introduce new artificial noise into the data. In contrast, STACCato incorporates batch effects as covariates in the tensor-based regression model, modeling batch effects jointly with other in-sample variables. That is, STACCato offers a comprehensive solution by jointly adjusting for multiple potential confounding variables, including both continuous (e.g., age) and categorical (e.g., batches, gender) variables.

Although both STACCato and Tensor-cell2cell utilize tensor-based techniques, their outputs and purposes are fundamentally different. STACCato operates on a CCC event-level, using a supervised regression model to directly assess the relationship between individual CCC events and the condition of interest, offering clear, actionable insights. In contrast, Tensor-cell2cell operates on a factor-level, using an unsupervised factor decomposition model that focuses on associations between decomposed factors and conditions, without providing explicit inference for specific CCC events. This distinction in output makes it difficult to directly compare Tensor-cell2cell with CCC event-focused methods like STACCato, which is better suited for targeted analyses and potential therapeutic interventions.

Compared to aggregation procedures, STACCato offers a major advantage by providing statistical inference (e.g., p-values) for associations between CCC events and conditions while jointly adjusting for multiple potential confounding variables. Moreover, the aggregation procedure is practically limited to categorical conditions, as it requires one aggregated sample per condition, making it ineffective for identifying CCC events related to continuous conditions like age. While the separate regression approach overcomes some limitations of aggregation procedures, our simulation studies and real-data analysis show that it is more sensitive to noise and outliers, leading to less reliable and more variable outcomes. In contrast, STACCato, by leveraging a tensor-based regression model, performs joint inferences across all CCC events, allowing information to be shared among correlated events. This makes STACCato practically superior, as it provides more stable and robust estimates of condition effects on CCC events, particularly in smaller datasets where noise and outliers have a greater impact, offering more reliable insights for follow-up analyses.

STACCato is a highly adaptable framework that can be seamlessly integrated with various existing CCC inference tools. Researchers have the flexibility to select any tool to calculate communication scores. For example, LIANA+ tool (Dimitrov *et al*., 2022, 2023) offers a comprehensive suite of tools and resources, such as CellPhoneDB (Efremova *et al*., 2020), CellChat (Jin *et al*., 2021), SingleCellSignalR (Cabello-Aguilar *et al*., 2020), to compute communication scores across all CCC events, as well as a practice protocol to construct the 4-dimensional communication score tensor (see Data and materials availability). STACCato can be subsequently applied to the 4-dimensional communication score tensor that is constructed by LIANA+ to detect condition-related CCC events. The tools implemented in LIANA+ can be categorized into magnitude-based and specificity-based methods. Magnitude-based methods measure the strength of communication, while specificity-based methods assess how specific a communication is to a given pair of sender and receiver cell types. The main advantage of specificity-based methods is that they avoid selecting common or background communications, often referred to as ‘housekeeping’ communications. STACCato tests for CCC events with significantly different communication scores across biological conditions of interest. Therefore, housekeeping CCC events, which remain consistent across different biological conditions, will not show a significant association with the biological condition in STACCato. Given the structure of the regression model in STACCato, where communication scores act as the response variable, we recommend using magnitude-based methods to calculate communication scores for STACCato, which is expected to provide more intuitive interpretations for the estimated effect coefficients of the condition variable on CCC events.

The STACCato framework does have its limitations. First, in scRNA-seq data, many genes may not be actively expressed in single cells, resulting in a significant proportion of zero values in the cell-cell communication score tensor. While STACCato can accept a sparse communication score tensor as input (e.g., with over 95% zeros in the tensor constructed using ASD and SLE datasets), the current framework does not explicitly model this data sparsity. A future extension of STACCato involving sparse tensor analysis, which imposes sparsity constraints on the ligand-receptor pairs, may inherently address this zero-inflation problem. Second, like other existing CCC inference tools, STACCato relies on a literature-curated ligand-receptor database, which limits the identification of condition-related CCC events to those previously documented. Extending STACCato to identify novel ligand-receptor pairs is part of our ongoing research but falls outside the scope of this work.

To enable the use of STACCato by the public, we provide an integrated tool (see Data and materials availability) to: (1) perform tensor-based regression to estimate the effects of the biological condition of interest on CCC events while adjusting for other covariates; (2) use the bootstrapping resampling method to assess the significance level of condition-related CCC events; (3) conduct downstream analyses, including comparing significant CCC events across cell types and visualizing CCC events that are significantly associated with the condition of interest. In conclusion, we present STACCato as a valuable tool to effectively identify condition-related CCC events using multi-sample multi-condition scRNA-seq data.

## Author contributions

QD conducted data analysis and drafted the manuscript; JY and MPE conceptualized and led the project and edited the manuscript.

## Data and materials availability

The processed data of the SLE dataset in h5ad format are available from https://www.ncbi.nlm.nih.gov/geo/query/acc.cgi?acc=GSE174188. The log2-transformed UMI counts of the ASD dataset are available from https://cells.ucsc.edu/autism/downloads.html. Source code for LIANA+ is available from https://github.com/saezlab/liana-py. The protocol of using LIANA+ and Tensor-cell2cell to build 4-D communication score is available from https://github.com/saezlab/ccc_protocols. Source code for Scanpy is available from https://github.com/scverse/scanpy. Source code for STACCato is available from https://github.com/daiqile96/STACCato.

