## Supplementary Materials for "Identifying condition-related cell-cell communication events using supervised tensor analysis"

Qile Dai *et al.*

**This PDF file includes:**

Supplementary Methods
Figs. S1 to S9
Tables S1 to S2

### Supplementary Methods

#### Construction of 4-dimensional communication score tensor

For a given scRNA-seq sample $i$ and a specific ligand-receptor pair $j$, we first compute its communication score across all $K$ sender cell types and $L$ receiver cell types and create a communication score matrix. In this matrix, the rows represent $K$ sender cell types; the columns represent $L$ receiver cell types; and the element located in the $k^{th}$ row and $l^{th}$ column corresponds to the value of $y_{ijkl}$. By repeating this process for all $J$ ligand-receptor pairs, we will get $J$ matrices, which can be arranged into a 3-dimensional tensor with dimensions $J \times K \times L$ for sample $i$. By repeating this process for all scRNA-seq samples, the 3-dimensional tensor of all $I$ samples can then be arranged into a 4-dimensional tensor $\mathcal{Y} \in \mathbb{R}^{I \times J \times K \times L}$. The 4 dimensions corresponds to $I$ samples, $J$ ligand-receptor pairs, $K$ sender cell types, and $L$ receiver cell types, with the $(i, j, k, l)$ entry of $\mathcal{Y}$ corresponds to $y_{ijkl}$. **Fig. 1B** shows a graphic example of building the 4-dimensional communication score tensor for a scRNA-seq dataset with three samples, two ligand-receptor pairs, and two sender-receiver cell types.

#### Rank selection by STACCato

We first determine the number of components $r_J$, $r_K$, $r_L$ for ligand-receptor pair, sender cell type, and receiver cell type dimension. For each dimension, we start by performing tensor unfolding to rearrange the elements of the communication score tensor into a matrix. For example, for the ligand-receptor pair dimension, we transform $\mathcal{Y} \in \mathbb{R}^{I \times J \times K \times L}$ into a matrix $Y_{(J)}$ with $J$ rows and $I \times K \times L$ columns. Then we set $r_J$ as the number of components that each can explain more than 1% of the variation in $Y_{(J)}$. We follow the same approach to determine $r_K$ for sender cell type dimension and $r_L$ for receiver cell type dimension. We set $r_Q$ as the number of sample-level variables available in $\boldsymbol{X}$.

**QR-adjusted optimization algorithm used by STACCato**

Denoting the supervised decomposition rank $\boldsymbol{r} = (r_Q\ ,\ r_J\ ,\ r_K\ ,\ r_L)$, we follow the optimization algorithm proposed by Hu et al. (Hu *et al.*, 2022) to estimate $\mathcal{B}, \mathcal{G}$, $\boldsymbol{M}_Q$, $\boldsymbol{M}_J$, $\boldsymbol{M}_K$ $\boldsymbol{M}_L$:

**Algorithm 1:**

Input: communication score tensor $\mathcal{Y} \in \mathbb{R}^{I \times J \times K \times L}$, sample-level design matrix $\boldsymbol{X} \in \mathbb{R}^{I\ \times Q}$, rank $\boldsymbol{r}$.

1. Normalize sample-level design matrix via QR factorization $\boldsymbol{X} = \boldsymbol{QR}$.
2. Project $\mathcal{Y}$ to the multilinear sample-level variable space to obtain the unconstrained coefficient tensor: $\widetilde{\mathcal{B}} = \mathcal{Y}\ \times_1 \boldsymbol{Q^T}$.
3. Obtain rank-constrained coefficient tensor by performing a rank- $\boldsymbol{r}$ higher-order orthogonal iteration (HOOI) (Kolda and Bader, 2009) on $\widetilde{\mathcal{B}}$: $\widehat{\mathcal{B}}^{(0)} \leftarrow HOOI(\widetilde{\mathcal{B}}, \boldsymbol{r})$.
4. Obtain estimated coefficient tensor by re-normalizing $\widehat{\mathcal{B}}^{(0)}$ back to the original feature scales: $\widehat{\mathcal{B}} = \widehat{\mathcal{B}}^{(0)} \times_1 \boldsymbol{R}^{-1}$.
5. Estimate $\mathcal{G}, \boldsymbol{M}_Q$, $\boldsymbol{M}_J$, $\boldsymbol{M}_K$, $\boldsymbol{M}_L$ by performing a rank-$\boldsymbol{r}$ HOOI on $\widehat{\mathcal{B}}$: $\widehat{\mathcal{B}} = \hat{\mathcal{G}}\ \times \{\widehat{\boldsymbol{M}_Q}, \widehat{\boldsymbol{M}_J}, \widehat{\boldsymbol{M}_K}, \widehat{\boldsymbol{M}_L}\}$.

Output: $\widehat{\mathcal{B}}$, $\hat{\mathcal{G}}$, $\widehat{\boldsymbol{M}_Q}$, $\widehat{\boldsymbol{M}_J}$, $\widehat{\boldsymbol{M}_K}$, $\widehat{\boldsymbol{M}_L}$.

We also impose orthonormality on $\boldsymbol{M}_Q$, $\boldsymbol{M}_J$, $\boldsymbol{M}_K$ $\boldsymbol{M}_L$ to ensure the uniqueness of decomposition.

**Parametric bootstrapping for hypothesis testing**

Denote the estimated communication score tensor as $\hat{\mathcal{Y}} = \widehat{\mathcal{B}}\ \times_1 \boldsymbol{X}$ with entry $\hat{y}_{ijkl}$ and the estimated standard error of $\epsilon_{ijkl}$ as $\hat{\sigma}$. We construct residual tensor $\mathcal{S} =\ \mathcal{Y} - \hat{\mathcal{Y}}$ with entry $s_{ijkl} =$

$y_{ijkl} - \hat{y}_{ijkl}$, and $\hat{\sigma}^2 = var(\text{vec}(\mathcal{S}))$, where $\text{vec}(\mathcal{S}) = [s_{1111}, \cdots, s_{ijkl}]$ denotes the vectorized version of tensor $\mathcal{S}$.

For the $n^{th}$ bootstrap resampling (Efron and Tibshirani, 1994), we generate a new tensor $\mathcal{S}^{(n)}$ with entries from $N(0, \hat{\sigma}^2)$ and construct a new communication score tensor $\mathcal{Y}^{(n)} = \hat{\mathcal{Y}} + \mathcal{S}^{(n)}$. We perform STACCato on $\mathcal{Y}^{(n)}$ to estimate a new coefficient tensor $\hat{\mathcal{B}}^{(n)}$. We repeat this procedure for $N$ iterations to generate $\hat{\mathcal{B}}^{(1)}$, $\hat{\mathcal{B}}^{(2)}$, $\cdots$, $\hat{\mathcal{B}}^{(N)}$. To test the null hypothesis of $H_0\colon \beta_{q,jkl} = 0$, we follow the guideline suggested by Hall and Wilson to define the bootstrap p-value as:

$$p_{q,jkl} = \frac{\sum_{n=1}^{N} I\left(\left|\hat{\beta}_{q,jkl}^{(n)} - \hat{\beta}_{q,jkl}\right| > |\hat{\beta}_{q,jkl}|\right)}{N+1}$$

where $\hat{\beta}_{q,jkl}^{(n)}$ denotes the $(q, j, k, l)$ entry of $\hat{\mathcal{B}}^{(n)}$; $\hat{\beta}_{q,jkl}$ denotes the $(q, j, k, l)$ entry of $\hat{\mathcal{B}}$, which is the estimated effect of variable $q$ on the CCC events involving the ligand-receptor pair $j$ between sender cell type $k$ and receiver cell type $l$; and $p_{q,jkl}$ is the bootstrapping p-value for $\hat{\beta}_{q,jkl}$.

**scRNA-seq dataset of SLE patients and controls**

The SLE scRNA-seq dataset collects multiplexed scRNA-seq of 264 PBMC samples from 162 SLE patients and 99 healthy controls (Thompson *et al.*, 2022; Perez *et al.*, 2022). The SLE scRNA-seq dataset in h5ad format was obtained from NCBI's Gene Expression Omnibus with GEO accession number 174188 (see Data and materials availability). From the h5ad data, we extracted the raw UMI counts of 32,738 genes across 1,263,676 cells from 264 samples and 99 technical replicates. We utilized Scanpy (see Data and materials availability) to filter out genes expressed in fewer than 3 cells, exclude cells with fewer than 200 expressed genes, normalize the raw UMI counts by count depth, and apply a log1p transformation. We then reduced the dataset down to one sample per subject by selecting the sample with the largest number of cells.

The metadata, which was also extracted from the h5ad data, includes the information of age, processing batch, ancestry, and gender of subjects. 107 (41%) subjects are Asian, 149 (57%) subjects are European, 3 (1%) subjects are African American, and 2 (1%) subjects are Hispanic. 13 (5%) subjects are flare SLE patients, 3 (1%) are treated SLE patients, 146 (56%) are managed SLE patients, and 99 (38%) are healthy controls. We filtered out 5 subjects of African American or Hispanic ancestry, as well as 13 flare SLE patients and 3 treated SLE patients.

The samples were collected from four processing cohorts. After filtering out samples from individuals of African American or Hispanic ancestry, as well as flare and treated patients, cohort 1 includes 42 healthy controls; cohort 2 includes 4 healthy controls and 105 managed SLE patients; cohort 3 includes 8 healthy controls and 1 managed SLE patient; and cohort 4 includes 44 healthy controls and 40 managed SLE patients. We only utilized the samples from cohort 2 and cohort 4 for our analysis, as cohort 1 consisted solely of healthy controls, and cohort 3 has only one managed SLE patient. In our analyses, we referred cohort 2 as Batch 1, and cohort 4 as Batch 2. We then utilized the processed expression data, along with the cell type information provided in the dataset's metadata, as input for LIANA+ to calculate communication scores and construct the communication score tensor for the remaining samples from 145 managed SLE patients and 48 healthy controls.

**scRNA-seq dataset of ASD patients and controls**

For the ASD dataset, we downloaded the raw UMI counts of PFC samples and the corresponding metadata from the UCSC Cell Browser (see Data and materials availability). This dataset contains the expression levels of 65,217 genes across 104,559 cells from 13 ASD patients and 10 healthy controls (Nassir *et al.*, 2021). We utilized Scanpy (see Data and materials availability) to filter out genes expressed in fewer than 3 cells, exclude cells with fewer than 200 expressed genes, normalize the raw UMI counts by count depth, and apply a log1p transformation. We then utilized the processed data, along with the cell type information provided in the dataset's metadata, as input for LIANA+ to construct the communication score tensor.

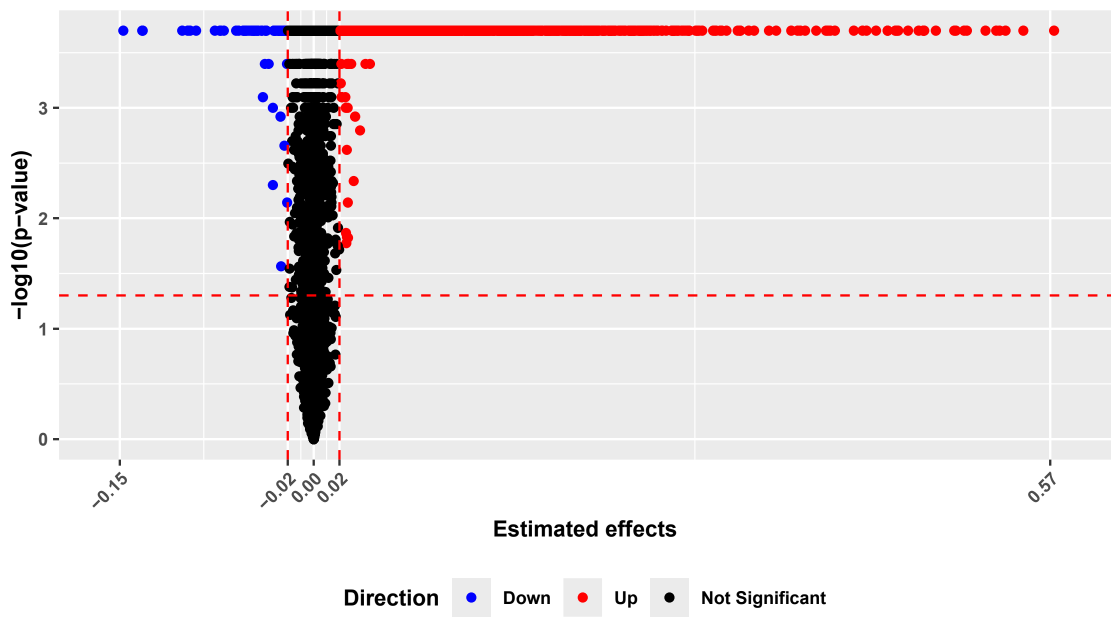


**Fig. S1. Significance levels of the estimated disease effects of all CCC events in the SLE dataset, generated by 4,999 iterations of bootstrapping resampling.** Significant SLE-related CCC events with p-values < 0.05 and magnitudes of their estimated disease effects > 0.02, are colored blue for negative effects and red for positive effects. Non-significant ones are shown in black. Y-axis represents -log10(p-values) and x-axis represents the estimated disease effects.

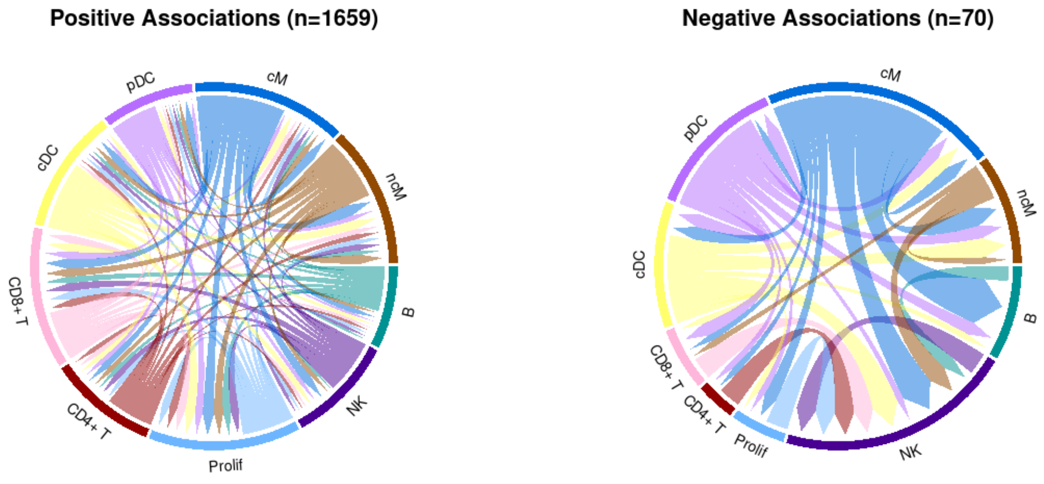


**Fig. S2. Circos plots where width of the links represents the total number of significant SLE-related CCC events between each pair of sender-receiver cell types.** A total of 1,659 CCC events demonstrate significant positive associations with SLE, while 70 CCC events demonstrate significant negative associations with SLE.

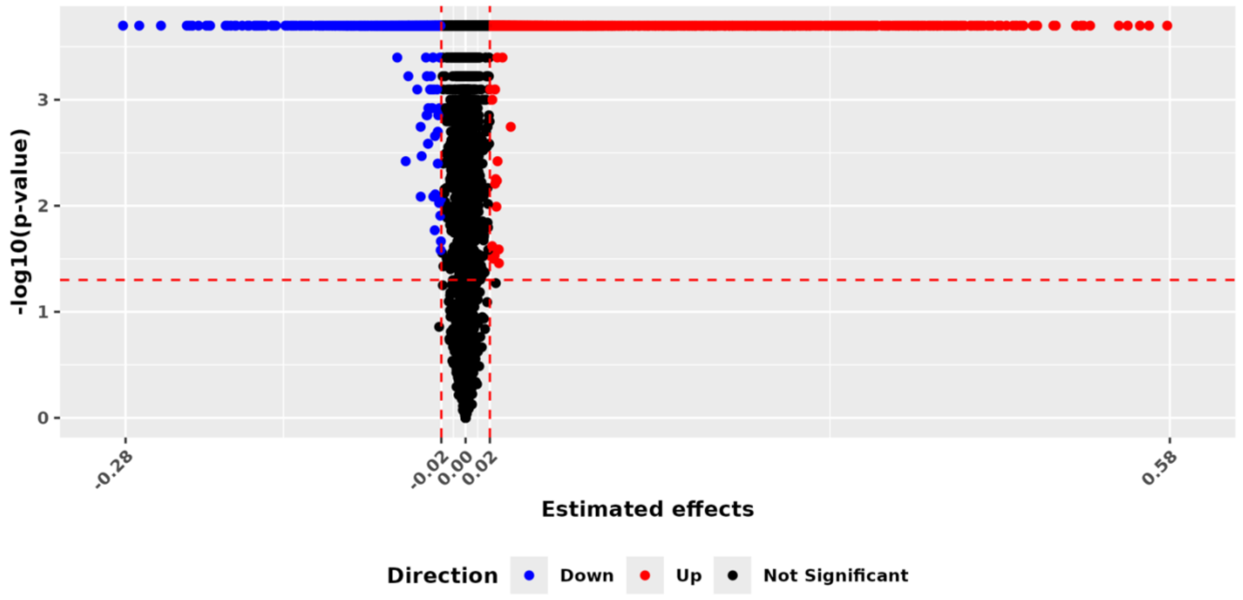


**Fig. S3. Significance levels of the estimated disease effects of all CCC events in the ASD dataset, generated by 4,999 iterations of bootstrapping resampling.** Significant ASD-related CCC events with p-values < 0.05 and magnitudes of their estimated disease effects > 0.02, are colored blue for negative effects and red for positive effects. Non-significant ones are shown in black. Y-axis represents -log10(p-values) and x-axis represents the estimated disease effects.

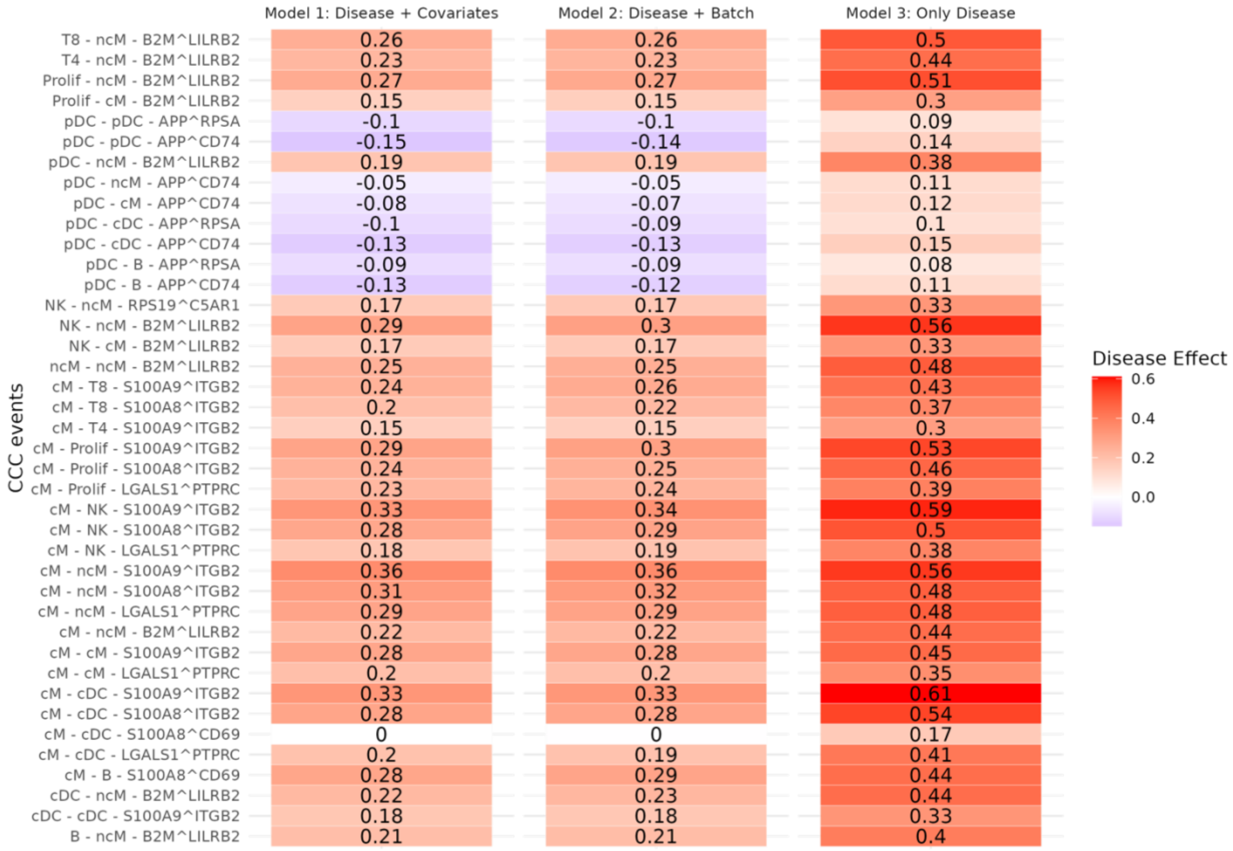


**Fig. S4. Comparison of the SLE effects estimated from three STACCato models with different sets of covariates.** Model 1 uses all available variables, including age, gender, ancestry, batch, and disease; Model 2 uses disease and batch variables; Model 3 uses disease variable only. Top 40 CCC events with the largest differences in estimated effects between Model 1 and Model 3 are plotted. The rows represent CCC events, formatted as sender cell type – receiver cell type – ligand^receptor pair. Positive disease effects are colored in red while negative disease effects are colored in blue.

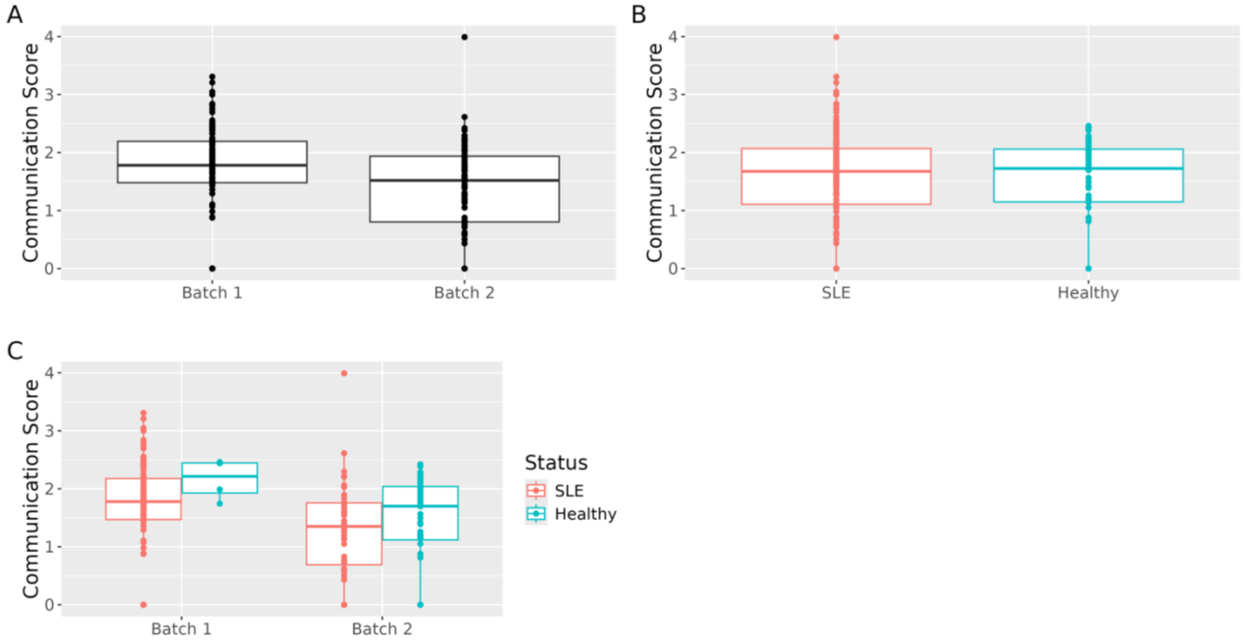


**Fig. S5. Box plots of the communication scores of the CCC event between APP and CD74 in pDC cells within the SLE dataset.** **(A)** Comparison between two batches; **(B)** Comparison between managed SLE patients and healthy controls; **(C)** Comparison between managed SLE patients and healthy controls within two batches.

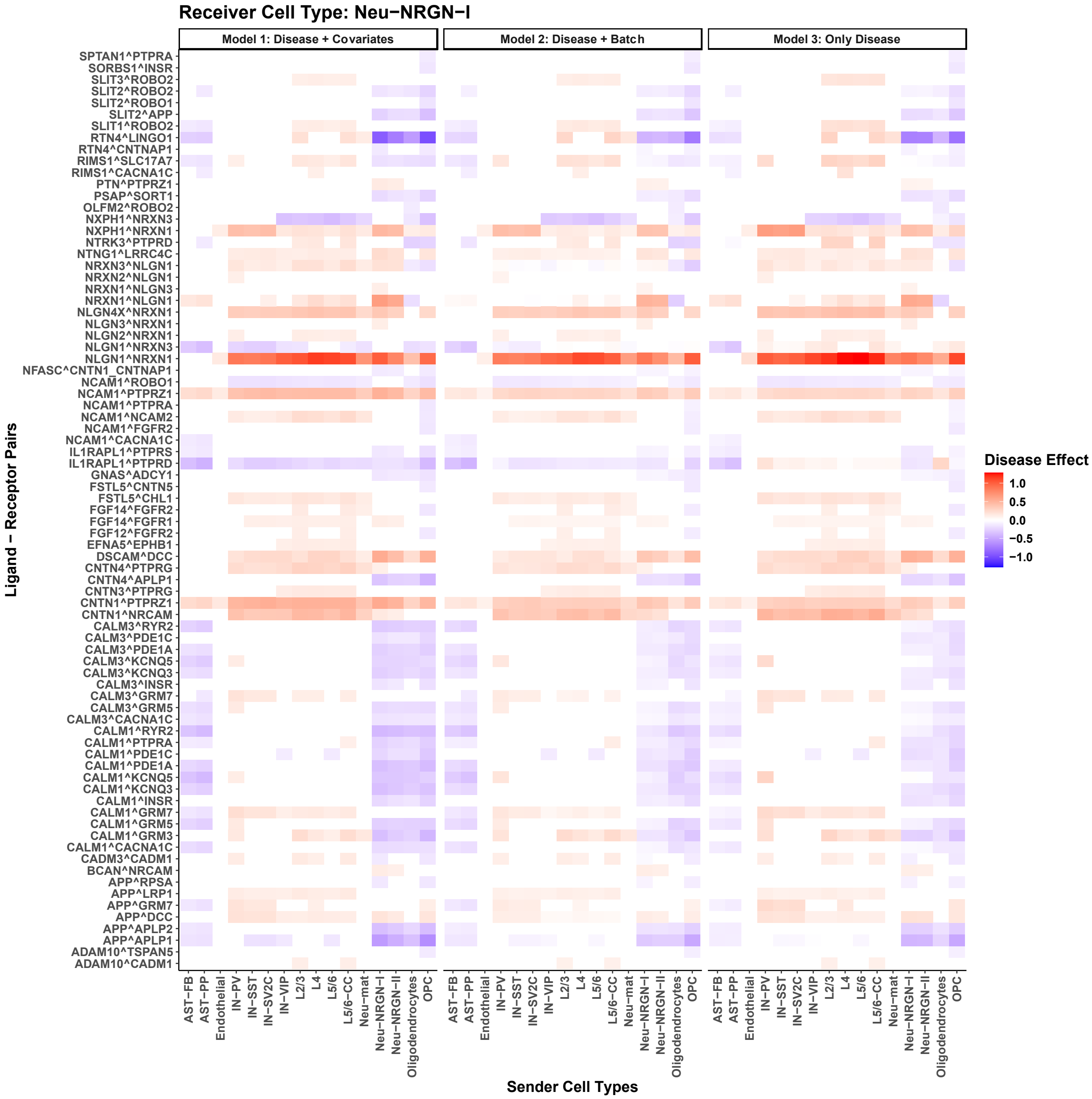


**Fig. S6. Comparison of the ASD disease effects for CCC events with Neu-NRGN-I as receiver cell type, estimated from three models with three different sets of covariates.** Model 1 uses all available variables, including age, gender, batch, and disease; Model 2 uses disease and batch information; Model 3 uses only disease. The disease effects of top 500 significant CCC events with Neu-NRGN-I as the receiver cell type by Model 1 are plotted. The rows represent ligand-receptor pairs, formatted as ligand^receptor. Positive disease effects are colored in red while negative disease effects are colored in blue.

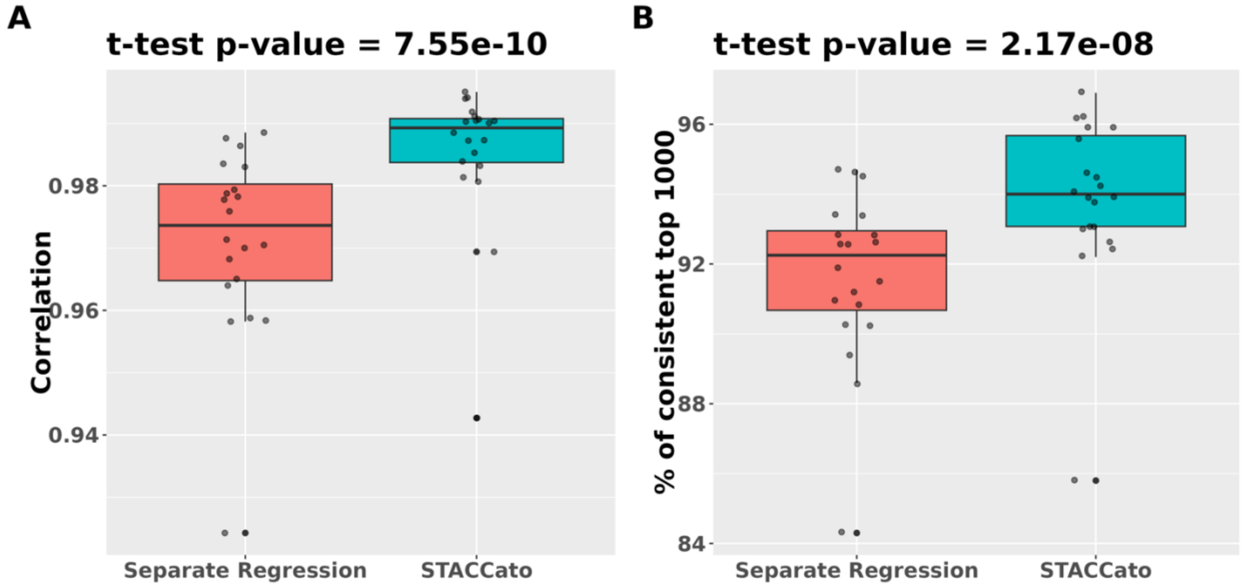


**Fig. S7. Robustness evaluations of STACCato and separate regression methods using the SLE dataset. (A).** The Pearson correlation coefficients of the estimated disease effects in 20 sub-datasets with those in the full dataset. Higher correlations mean more robustness; **(B).** The proportion of the top 1000 significant CCC events (those with the largest magnitudes of disease effects) in the full dataset that also appear in the 20 sub-datasets. Higher proportions mean more robustness. We used a paired t-test to determine whether the Pearson correlations and overlap proportions were significantly different between STACCato and the separate regression approach. T-test p-values are significant for both comparisons and are shown in the figure.

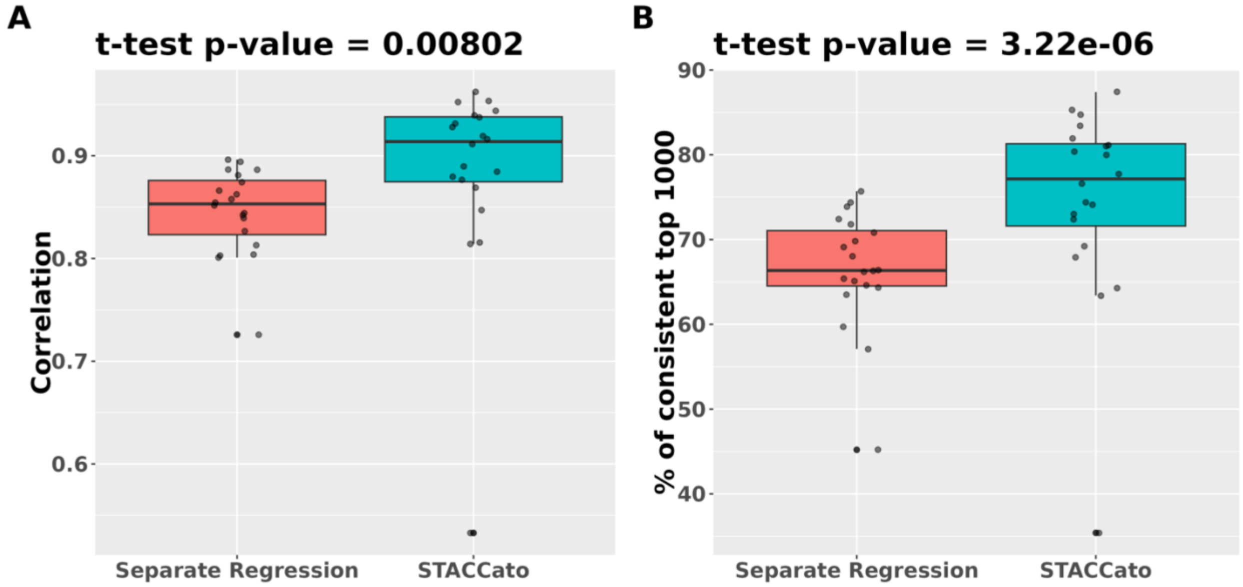


**Fig. S8. Robustness evaluations of STACCato and separate regression methods using the ASD dataset. (A).** The Pearson correlation coefficients of the estimated disease effects in 20 sub-datasets with those in the full dataset. Higher correlations mean more robustness; **(B).** The proportion of the top 1000 significant CCC events (those with the largest magnitudes of disease effects) in the full dataset that also appear in the 20 sub-datasets. Higher proportions mean more robustness. We used a paired t-test to determine whether the Pearson correlations and overlap proportions were significantly different between STACCato and the separate regression approach. T-test p-values are significant for both comparisons and are shown in the figure.

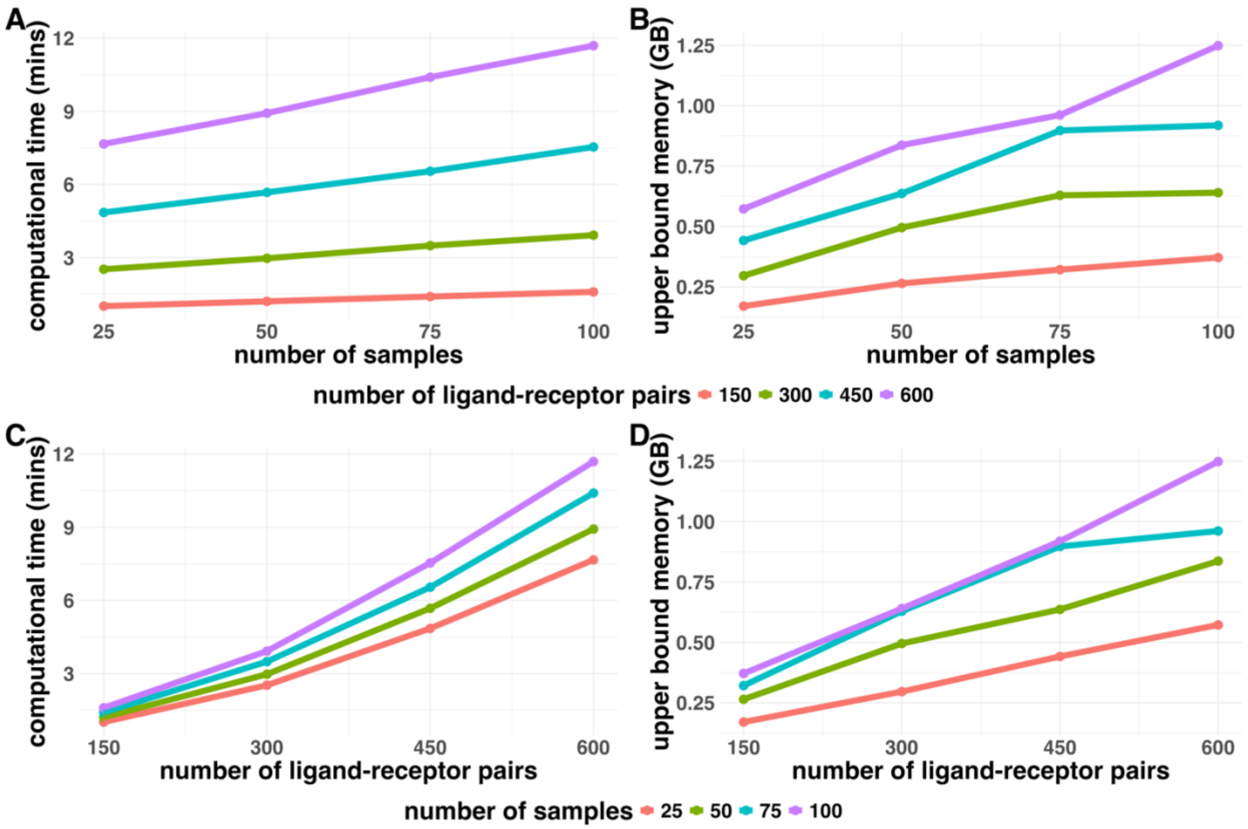


**Fig. S9. Computation CPU time and memory usage of 99 iterations of bootstrapping resampling in simulated datasets with various numbers of samples and ligand-receptor pairs.** The simulated datasets are of 10 sender and receiver cell types and 10 sample-level variables. **(A, B).** Y-axis shows the computational CPU times are in minutes (mins) and upper bounds of memory usages are in gigabytes (GB), with various numbers of samples $\boldsymbol{I} = (\mathbf{25}, \mathbf{50}, \mathbf{75}, \mathbf{100})$ in the x-axis and various numbers of ligand-receptor pairs $\boldsymbol{J} = (\mathbf{150}, \mathbf{300}, \mathbf{450}, \mathbf{600})$ in different colored lines. **(C, D).** Y-axis shows the computational CPU times and upper bounds of memory usages, with various numbers of ligand-receptor pairs in the x-axis and various numbers of samples in different colored lines.

**Table S1. The number of healthy controls and managed SLE patients in each batch of the SLE dataset.**

| | Batch 1 | Batch 2 |
|---|---|---|
| Healthy controls | 4 | 44 |
| Managed SLE patients | 105 | 40 |

**Table S2. The number of healthy controls and ASD patients in each batch of the ASD dataset.**

| | Batch 1 | Batch 2 |
|---|---|---|
| Healthy controls | 5 | 5 |
| ASD patients | 5 | 8 |
